# Adaptation to increasing elevation via metabolic rates and mitochondrial genetics in Andean birds

**DOI:** 10.64898/2026.10.12.758415

**Authors:** Erik N. K. Iverson, David Guevara-Apaza, Erick Huaman Vargas, Ash Romero, Emily A. Pearson, Beril L. Saygin, Daniela A. Florez Barra, Daira A. Santos Valencia, Justin C. Havird

**Affiliations:** Department of Integrative Biology, University of Texas at Austin, Austin, TX; Wildlands Studies, Aptos, CA; Department of Bioengineering, University of Pennsylvania, Philadelphia, PA; Facultad de Ciencias Biológicas, Universidad Nacional de San Antonio Abad de Cusco, Cusco, Perú; Department of Entomology, University of California, Davis, Davis, CA; Dell Medical School, University of Texas at Austin, Austin, TX

## Abstract

Closely related species often replace each other at different elevations, a pattern that results in high biodiversity in mountains. Species range limits are set by physiological adaptation to gradients in temperature and oxygen availability as well as by competition with species above and below them on the gradient. As mountains warm due to climate change and many species move upward in elevation it is increasingly important to understand the mechanisms of climatic adaptation and competition that govern where and when species will persist, migrate, or go extinct. We investigated the genetics and physiology of tapaculos (Passeriformes: Rhinocryptidae), a group of small, weakly flying birds known for their narrow elevation ranges across the Andes, where as many as 6 species may replace each other on one transect. We first looked for selection on the mitochondrial genome, which is thought to be important in adaptation to elevation, in 85 members of this clade across South America. We found a signature of intensified positive selection on mitochondrial genes among species with mid-point elevations above 3000 m, suggesting metabolic adaptation to the harsh, high-altitude environment. We then measured metabolic rates and mitochondrial function in the field in four species of the genus *Scytalopus* that replace each other in a series along the Manu elevation gradient in Southeastern Peru. We found that higher elevation species have higher Basal Metabolic Rates (BMR) and more extreme Lower Critical Temperatures (LCTs), suggesting adaptation to the cold. However, these differences do not have a clear basis in mitochondrial function, which was broadly similar across species. Rather, higher-elevation species are smaller overall but have larger relative organ masses and maintain lower body temperatures, which may be adaptations to meeting the high thermogenic demands of their environment. These four species conform well to expectations of the Competitive-Exclusion Tolerance Rule (Martin & Ghalambor, 2023), with each species up the gradient being a poorer competitor (smaller body size) but having adapted to a harsher part of the gradient that more competitive species cannot occupy. We discuss these results in the context of theories about the importance of climate, competition, and ecotones in the distribution of Andean organisms, predicting that species living in the highest-elevation forests just below the treeline may be at the greatest risk of extinction due to climate change.

## Introduction

Tropical mountains have long served as natural ecological and evolutionary laboratories because they present an extremely diverse set of climates in a relatively small geographic area (Feeley et al., 2011; Humboldt & Bonplan, 1807; Janzen, 1967; Mayr & Gilliard, 1954; Terborgh, 1971). Mountains recapitulate the climatic zonation seen moving from the tropics to the poles, and many taxa show phylogenetic segregation and ecological replacement along elevational gradients that mirrors latitudinal replacement. Elevation ranges in related tropical taxa, for instance, often occur in discrete, narrow bands of stable climate along mountains (Janzen, 1967; McCain, 2009). It is unclear whether speciation is a consequence or a cause of climatic specialization, and which genes and physiological constraints are involved (Cadena et al., 2020; Keller & Seehausen, 2012; Linck et al., 2020). However, species replacing each other on an ecological gradient are rarely thought to have formed *in situ* through parapatric speciation; rather, current opinion holds that they typically diverge in a period of isolation (Cadena & Céspedes, 2020; Caro et al., 2013; Linck et al., 2020; Rossetto et al., 2012), and, upon secondary contact, the boundary between species is established by the interplay of competition and environmental tolerance (Martin & Ghalambor, 2023). Understanding how climatic niche segregation evolves and what processes set the environmental boundaries between species will be essential to predicting how and why biodiversity will be redistributed by climate change.

On mountain gradients, metabolic phenotypes, driven by interactions between mitochondrial and nuclear genes, might be one of the primary traits that diverge in allopatry and create reproductive isolation between species. Studies frequently report signatures of selection in high-elevation populations and species in both the mitochondrial genome (Graham et al., 2024; Jin et al., 2021; Li et al., 2018) and in nuclear genes involved in energy metabolism, angiogenesis, and other key pathways (Bigham, 2016; Jacovas et al., 2018; Tiwari et al., 2024). Some of the adaptive differences between low- and high-elevation taxa are attributable to evolved and plastic mitochondrial responses to cold (reviewed in Scott et al., 2018), including differences in the activity of mitochondrial enzymes (Garrett et al., 2024) and uncoupling of the mitochondrial respiratory chain and increased leak in highlanders, which generates heat (Mahalingam et al., 2020). In endotherms, which have a highly aerobic metabolism that sustains an elevated internal body temperature, high-elevation species face selection both to produce more heat and to do so with less available oxygen, leading to differences in metabolic and mitochondrial processes (Cheviron et al., 2012; Gutierrez-Pinto et al., 2021; Rosenmann & Morrison, 1975). For instance, within deer mice (*Peromyscus maniculatus*), high-altitude populations display evolved differences in thermogenic capacity, including lower metabolic rates under hypoxic conditions, demonstrating adaptation to a low-oxygen environment (Bautista et al., 2024; Cheviron et al., 2012, 2013, 2014; Tate et al., 2020).

With high body temperatures and an energetically demanding lifestyle, birds are some of the organisms that might face strong selection on metabolic performance to adapt to different elevations. For instance, extreme high-altitude birds show a variety of molecular and vascular adaptations to low-oxygen conditions (reviewed in Laguë, 2017). However, it is unclear whether elevated metabolic rates are a general feature of avian adaptation to altitude, as might be expected for endotherms. Higher basal metabolic rates (BMR) were not reported among high-elevation birds in Peru or New Guinea (Londoño et al., 2015; McNab, 2013), but a controlled comparison of congeners at different elevations in Colombia did find a pattern of increased BMR within genera (Gutierrez-Pinto et al., 2021). Like many tropical organisms, tropical bird genera show a high degree of replacement across elevations, with closely related species occupying narrow ranges at progressively higher elevations (Cadena et al., 2020; Cadena & Céspedes, 2020; Terborgh, 1971). Due to a lack of seasonality, tropical organisms are expected to adapt to the narrow set of conditions present within their range (Janzen, 1967; McCain, 2009).

The question of to what degree species have adapted to their particular elevation ranges is relevant because birds, like many other tropical organisms, are moving upward in elevation due to rising temperatures (Feeley et al., 2011; Freeman et al., 2018; Freeman & Class Freeman, 2014; Morueta-Holme et al., 2015; Neate-Clegg et al., 2021), potentially causing an “escalator to extinction” on mountains of limited height (Freeman et al., 2018; Urban, 2018). However, it is not clear which processes are driving this redistribution. Related birds might have broadly similar environmental tolerances across elevations and be responding to shifting patterns in food and habitat; alternatively, if birds are well-adapted only to their particular climates, they may be tracking temperatures directly. It is also important to understand how environmental tolerance interacts with competition from neighboring congeners to set and shift range limits (Freeman et al., 2022; Martin & Ghalambor, 2023).

The small neotropical birds known as *tapaculos* (Passeriformes: Rhinocryptidae) are ideally suited to address questions of adaptation and specialization on elevation gradients, being found from sea-level to montane grasslands over 4000 m high across the Andes (Cadena et al., 2020; Cadena & Céspedes, 2020). Their expansion has been deemed a ‘non-adaptive radiation’ as, for instance, the 45 species in the genus *Scytalopus* have retained a similar appearance and behavior across widely differing environments (Cadena et al., 2020). However, they almost certainly have adapted in their metabolic and thermal traits during this radiation, as their narrow ranges vary by up to 20 °C in average temperature. Along a single elevational transect, as many as six species may replace each other with increasing elevation (Cadena & Céspedes, 2020). Adjacent taxa on the gradient are typically not each other’s closest relatives, supporting a model of allopatric speciation followed by secondary contact (Cadena et al., 2020; Cadena & Céspedes, 2020; Linck et al., 2020). In order to examine patterns of environmental selection and ecological specialization in this group, we characterized the signature of both positive and relaxed selection across the 13 mitochondrial protein-coding genes in 85 tapaculo taxa in relation to their elevation and latitudinal range limits and range sizes. We then captured members of four *Scytalopus* species that replace each other with elevation along a Peruvian transect and measured whole-animal and mitochondrial metabolic rates in the field, predicting that higher-elevation species would have higher basal and summit metabolic rates underlain by differences in mitochondrial function. Our analyses shed light on the nature of endotherm environmental adaptation and inform predictions about how these species will respond to warming across elevations.

## Methods

### Selection on Mitochondrial DNA Across Elevation in Tapaculos

#### Data curation

To analyze selection on metabolic genes in a phylogenetic context, we examined the 13 mt protein-coding genes. Our approach was similar to that in Iverson et al. (2025), quantifying two nonsynonymous substitution patterns to differentiate positive from relaxed selection. Ultra-conserved element (UCE) data are available for almost all members of the speciose tapaculo genus *Scytalopus* (Cadena et al., 2020) and several other species in Rhinocryptidae, as well as the sister family Melanopareidae (cresent-chests). We first extracted the mt CDS from all available species in these two families using MitoFinder (Allio et al., 2020). MitoFinder was able to recover one or more genes from 68 samples of *Scytalopus* (including several conspecific samples in some cases), 19 samples from other tapaculo genera (Rhinocryptidae), and 5 species from Melanopareidae which were used as outgroups. These 92 taxa, for which we extracted from 1 to 13 mt genes, were used to construct a bayesian phylogeny of the entire group. Additionally, sequences of *ND2* or *COX1* from NCBI’s *nuccore* were used to add three additional species to the phylogeny, as well as to augment the sequence of *S. intermedius* (SRR12463120) for which MitoFinder only recovered one gene. Each gene was aligned for all species together in MEGA (Kumar et al., 2018) using MUSCLE (Edgar, 2004) based on amino acid sequences. Stop codons were removed, and alignments were visually inspected and manually re-aligned where necessary. Alignments were concatenated into one 13-gene sequence (See Data Availability).

#### Phylogenetic analyses

We created four Bayesian phylogenies of the Rhinocryptidae with *Melanopareia* as the outgroup using MrBayes v. 3.2 (Ronquist et al., 2012). We used two different partition schemes and created phylogenies under each for both the 95 taxa available as well as the 85 taxa for which more than 5 protein-coding genes could be recovered. The first partition treated each of the 13 protein-coding genes as evolving under separate rates. The second partition gave each of the three codon positions in the entire concatenated sequence a distinct rate. All partitions were set to nst = 6 and rates = gamma with statefreqprset = dirichlet(1.0,1.0,1.0,1.0). Two MCMC runs with four chains ran for 6,000,000 generations at a temperature of 0.10 with sampling every 100 trees. MrBayes was run on the CIPRES Science Gateway web server (Miller et al., 2010) using BEAGLE (Ayres et al., 2012). Traces and parameter distributions were checked in Tracer v. 1.7.1 (Rambaut et al., 2018) and consensus trees were visualized using FigTree v. 1.4.4 (http://tree.bio.ed.ac.uk/software/figtree/). In initial runs, one taxon stood out as a branch length outlier, *S. magellanicus* (SRR12555200), and was placed as basal to all other species in the phylogeny. The alignment of this sequence was found to be correct, but it stood out for its high incidence of nonsynonymous substitutions. We concluded that MitoFinder may have failed to properly filter “NuMts” and that the sequence used was likely partly or entirely composed of these pseudogenized nuclear genes of mitochondrial origin. We therefore replaced this taxon with an incomplete CDS for the species (KJ909189).

#### Range-wide environmental parameters

Elevational range limits for each species in the phylogeny were obtained from either the IUCN or Cornell University and del Hoyo *et al*.’s *Birds of the World* (https://birdsoftheworld.org/bow/home). Latitudinal range limits and range sizes were extracted from IUCN shapefiles using a Python script from Iverson et al. (2025). Range size was included in the analysis because, as a proxy for effective population size (*N_e_*), small-ranged species may be more likely to show elevated *d*_N_/*d*_S_ due to relaxed selection (Iverson et al., 2025; see below). The midpoint of both elevational and latitudinal ranges was calculated, as well as a climatic index (see Iverson et al., 2025) expressing the relationship of both midpoints to their theoretical maximum. The climatic index was calculated as follows:

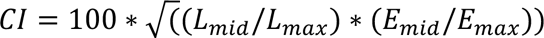

where *L_mid_* and *E_mid_* refer to the latitudinal and elevational midpoints of a given species, and *L_max_* and *E_max_* refer to the maximum values of elevation and latitude obtained by any species in the group.

#### Tests for selection

The *d*_N_/*d*_S_ ratio, reflecting nonsynonymous substitution bias, is a common metric of selection strength that has been used to support many claims of high-altitude adaptation (Iverson et al., 2025). However, *d*_N_/*d*_S_ ratios cannot distinguish between positive and relaxed selection. To draw conclusions about adaptation it is necessary to combine these metrics with explicit tests for relaxation of selection (see below). We first used the codeML model in the program PAML 4.0 (Yang, 2007) to calculate *d*_N_/*d*_S_ values among branches in the 85-taxon tree generated by the *position*-partition. The 85-taxon tree was used to limit selection tests to sequences with sufficient information, in those with more than 5 protein-coding genes out of 13. We ran a multi-rate model in codeML using default parameters for the vertebrate mitochondrial code and assigning tips of the phylogeny to different bins based on the ecological parameters above. Sequences from the same taxonomic species that clustered monophyletically were treated as one species unit, and the internal branches of such clusters were included in the same rate bin as the tips, whereas conspecific sequences that clustered with other species in the phylogeny were treated separately. All remaining internal branches of the phylogeny comprised the “background” bin for purposes of comparing tip rates. Thus, with one codeML run, we calculated separate *d*_N_/*d*_S_ values for each bin of tips and a background value (all internal branches).

We then used the RELAX web server (https://www.datamonkey.org/relax) to distinguish positive from relaxed selection, both of which can cause elevated *d*_N_/*d*_S_ values. RELAX (Wertheim et al., 2015) calculates a parameter, *k*, as a measure of the intensity of selection on a set of test branches relative to reference branches, with *k* > 1 indicating intensified selection (either positive or purifying) and *k* < 1 relaxed selection (which can also be either positive or purifying). RELAX then tests for the significance of the deviation in *k* from expected values. RELAX examines the distribution of *d*_N_/*d*_S_ values among sites and tests for a characteristic pattern of relaxed selection, where small *d*_N_/*d*_S_ values increase towards 1 while larger values decrease towards 1 in test branches relative to reference branches. Under positive selection, by contrast, all values move away from neutrality, with small values decreasing and large values increasing.

We used four different binning schemes to examine how *d*_N_/*d*_S_ and *k*-values changed across ecological gradients. First, we binned species by midpoint elevation, into 8 bins of 500 meters each (i.e., 0-499 m, 500-999 m, 1000-1499 m, etc.). Then we binned species by midpoint latitude, into 8 bins of 5° latitude. We then used the Climatic Index to bin species, into 10 bins of width CI = 5, beginning with 5-9.9. Finally, we binned species according to the natural log of their range size, into 6 bins of two log units (ln(range size) = 5-6, 7-8, 9-10, etc.). We dropped bins from our analysis if the *d*_N_/*d*_S_ that codeML estimated for that bin was 0.0001 or 999, indicating failure for the model to converge on a reliable estimate due to an insufficient number of substitutions. We ran an additional null model, where codeML calculated one *d*_N_/*d*_S_ rate for the entire phylogeny, to perform a likelihood ratio test for each binned model. We used the same branch partitioning schemes as for *d*_N_/*d*_S_ ratios, partitioning based on elevation, latitude, *CI*, or range size. RELAX was run once for each bin of each partition, with the tips in that bin as the test branches and the common set of internal branches (excluding all other tips) as the reference branches. Thus, each bin of tips was tested against common internal branches, and the *d*_N_/*d*_S_ values calculated above correspond to each of these bins and the background *d*_N_/*d*_S_ to these same internal branches.

### Metabolic Rates and Mitochondrial Physiology in Four Species that Replace Each Other with Elevation

#### An elevation transect in Peru as a natural laboratory

We examined four species of *Scytalopus* that replace each other with elevation in the Manu Biosphere Reserve of Southeastern Peru (Fig. 2.1). The species were, from lowest to highest elevation, *S. atratus* (800 – 3100 m, reported range-wide elevations), which occurs in montane rainforest, *S. parvirostris* (1630 – 4090 m), which occurs largely in cloud forest, *S. schulenbergi* (1760 – 4230 m), which occurs in upper cloud forest and elfin forest, and *S. simonsi* (2660 – 4660 m), which occurs in high-altitude *puna* grasslands above the treeline. In the area where we worked, the species occur, in order, from roughly 700 – 2500 m, 2000 – 3300 m, 3100 – 3700 m, and 3300 – 4000 m, with elevational overlap where habitat types intergrade. *S. atratus* were captured along the Manu Road (CU-113) and in the surroundings of the Cock-of-the-Rock Lodge at San Pedro (−13.05487, - 71.54668; 1380 m) in the buffer zone of Manu National Park and measured at the same elevation. *S. parivostris* were captured higher along the road and in the surroundings of Wayqecha Cloud Forest Biological Station (−13.17538, −71.58651; 2900 m) and measured at the biological station. *S. schulenbergi* and *S. simonsi* were captured inside Manu National Park’s Tres Cruces sector and measured at a common elevation inside a cabin located at the ridgetop (−13.12065, −71.61160; 3640 m). Birds were captured in mist nets using targeted playback between 14:00 and 18:00 and held with access to water but without food, in a covered transport cage away from disturbance. All procedures were carried out in compliance with IACUC protocols AUP 2019-00287 and AUP 2023-00015, between July and September 2022 and July and August 2023, and with relevant Peruvian permits (SERFOR: Expediente No. 2023-0022717; SERNANP: *Resolucion Jefatural del Parque Nacional del Manu* No. 13-2022-SERNANP-JEF).

**Figure 2.1:**
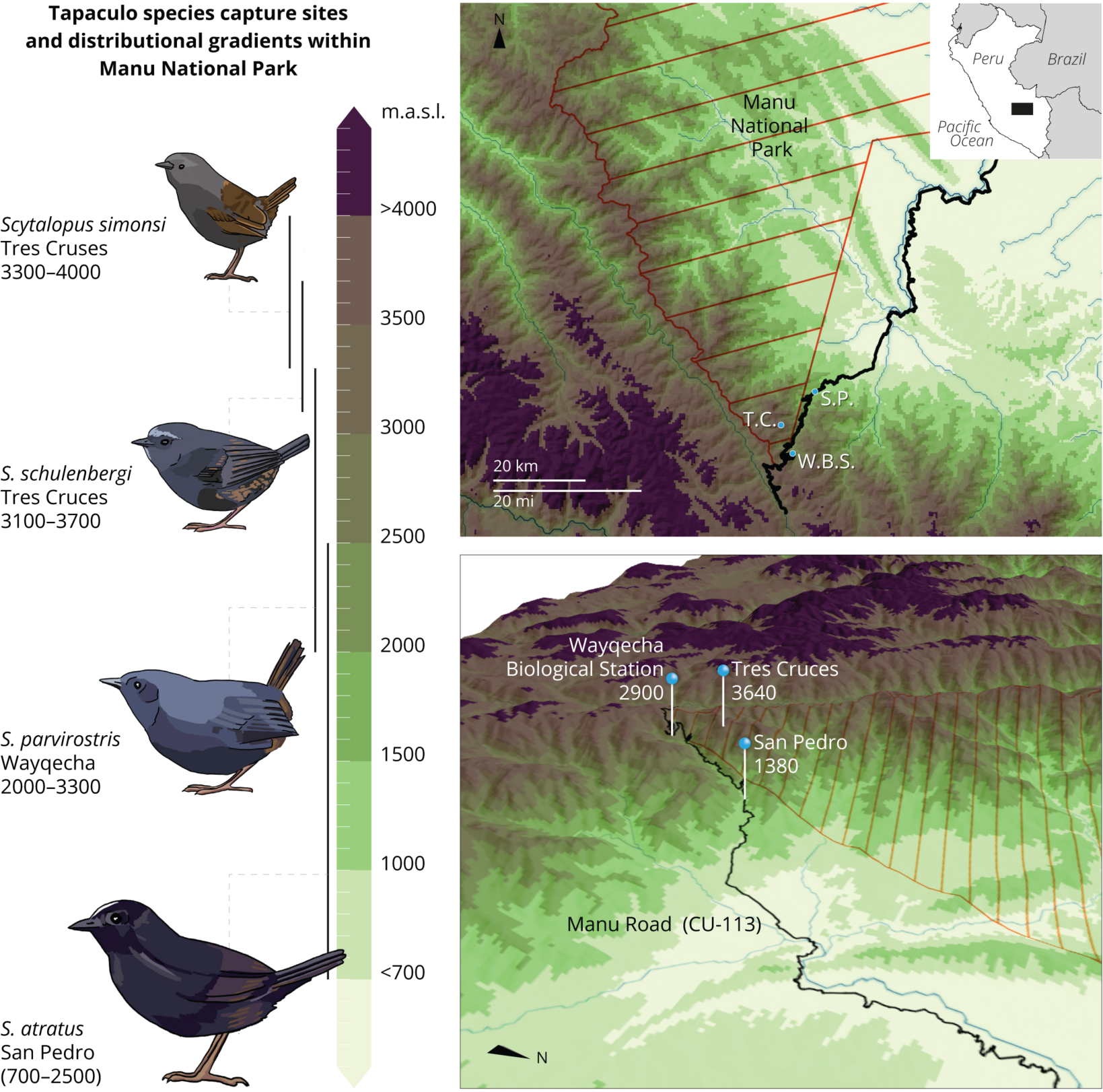
Location in the Manu Biosphere Reserve of Southeastern Peru, where four species of *Scytalopus* (*lefthand side*) were caught in 2022 and 2023. *S. atratus* were captured and measured at the Cock-of-the-Rock Lodge in San Pedro (elevation: 1380 m), *S. parvirostris* were captured and measured at Wayqecha Cloud Forest Biological Station (2900 m), and *S. schulenbergi* and *S. simonsi* were captured and measured at the cabin at Tres Cruces inside of Manu National Park (3640 m). *Upper panel*: overhead digital elevation model with colors corresponding to elevations on the lefthand scale bar; *lower panel*: angled view with labeled study locations. Black bars near each species represent the approximate elevation range of each species on the Manu elevation gradient. The four species are depicted at different sizes in accordance with their observed differences in mass (Fig. 2.4c). Cartography & figure design: Andrés Alegría. Tapaculo drawings: Emily Underwood.

#### Ambient temperature monitoring

To quantify differences in climatic niches between species that may influence adaptation, we used HOBO Pendant MX (Onset Computer Corporation, Bourne, MA) & Elitech RC-51H (Elitech, San Jose, California) temperature loggers to measured daily temperature and humidity patterns during a common two-week period in July-August 2023. We placed four loggers in the habitat type corresponding to each species (montane forest at 1380 m, cloud forest at 2900m, elfin forest and puna grasslands at 3640 m), in pairs covering two distinct microhabitats at two different locations. One logger was hung from a branch in the air 2 m from the ground, representing ambient conditions, and other was placed at ground level inside of dense vegetation of the kind commonly used by *Scytalopus* tapaculos. Loggers recorded every minute, and we calculated daily minimum and maximum temperatures across a common 17-day period (31 July to 6 Aug and 8 Aug to 17 Aug, 2023). While loggers were typically shaded by canopy cover and nearly continuous cloud cover, records with maximum temperature values above 30 °C were occasionally recorded at all elevations and were removed as erroneous values caused by the impact of direct sunlight on loggers. Values were averaged for replicate loggers in the same position and same habitat type.

#### Whole organism metabolic rates across decreasing temperatures

We quantified basal metabolic rate (BMR) by measuring fluxes of O_2_, CO_2_, and water vapor in a flow-through protocol across a decreasing temperature gradient using a Field Metabolic System from Sable Systems (Las Vegas, NV). Because of the need for slow ramping rates, we only had time to take birds through a protocol measuring either Upper Critical Temperature (UCT) or Lower Critical Temperature (LCT), which are the points bounding the thermoneutral zone of an endotherm. Between these temperatures, metabolic rate is constant, while above UCT or below LCT, endotherms increase metabolic rate to engage in active thermoregulation. While measuring UCT might appear more valuable for studying the impacts of climate change, we chose LCT as a better indicator of climatic specialization that might inform species’ responses to climate change. This was because UCTs commonly reported for tropical birds (34-40 °C) are well above the highest temperatures experienced at any elevation in our study area. By contrast, both daytime and nighttime temperatures in our study area (see Fig. 2.4A, B) frequently fall below reported LCTs for tropical birds (22-30 °C) (Monge et al., 2023; Pollock et al., 2021). Daytime highs are also much more similar across elevations than nighttime lows (Fig. 2.4A, B). We thus considered it unlikely that UCTs are under strongly differing selection between these species.

Trials for the measurement of metabolic rate began after sunset (18:30) and a minimum of 4 hr after capture, to ensure that birds were post-absorptive. Notably, we could not control for ambient air pressure in our metabolic rate trials; all birds were measured at native elevations under ambient air pressure. The Field Metabolic System recorded air pressure and this was used to correct and standardize readings across elevations. Before birds entered the experiment, we recorded cloacal temperature and body mass. Birds were placed into an inner chamber with incurrent and excurrent airflows, which was placed into a larger temperature-controlling cabinet. A long-stem thermometer enabled the observer to confirm temperature inside the inner chamber, which reflected the temperature settings of the external cabinet but with greater variability due to the movement, behavior, and thermoregulatory state of the bird.

The temperature of the cabinet was manually lowered over the course of the night in a standardized program, with the set temperature, realized temperature, and internal chamber temperature recorded every 10 minutes. Because ambient starting temperature may already have been lower than the LCT for some species or individuals, we began the experiment by first raising temperature to 28 C, which we considered likely to be above the LCT of a montane tropical bird (Monge et al., 2023), from which point it would be lowered past the LCT. The experiment began at ambient temperature and the temperature was raised by 0.1 degree / minute for 30-minutes; thereafter temperature was raised by 0.2 degrees / minute to 28 C, which was held for 30 minutes. Temperature was then lowered by 0.2 degrees/minute with 30-minute holds at 22 °C and 16 °C; below 12 °C, temperature was lowered at 0.1 degree / minute, with 30-minute holds at 10 °C and 4 °C. Below 4 °C, temperature was lowered at 0.05 degrees / minute, with a 30-minute hold at −2 °C. Cabinet temperatures below 0 °C were not possible at the low elevations where *S. atratus* was studied due to high ambient temperature, so metabolic rate was only measured at these temperatures for the other three species. A trial ended when the lowest temperature possible by the Field Metabolic System was reached, −5 °C was reached, or the bird exhibited signs of distress (rapidly elevating/fluctuating CO_2_ flux). Temperature was then raised from the stop point to the ambient temperature at the fastest rate the system would allow, approximately 1 degree / minute. In total, trials lasted 6-9 hrs in duration, and ended between 3:00 and 6:00 in the morning.

Metabolic rate was assessed continually in 3-minute windows where the system analyzed excurrent air from the chamber containing the birds, separated by 2-minute baseline windows (to control for sensor drift) analyzing fresh air that had passed through a length of tubing inside the temperature cabinet. These values were used to estimate LCT, while metabolic rate at defined hold periods (28, 22, 16, 10, 4, 2, 0, & −4 °C) was assessed by averaging the VO_2_ values for all windows where the measured internal chamber temperature (averaged over the window) was within 0.5 °C of the target hold temperature. VO_2_ measurements were cleaned and filtered with a custom macro in ExpeData.

We determined the LCT of VO_2_ (oxygen consumption rate) by performing a breakpoint analysis on cleaned and filtered VO_2_ measurements. We used the R package ‘segmented’ (v. 2.1) to calculate the AIC values of linear models with multiple slopes at different user-specified numbers of breakpoints and evaluated the breakpoints found for the model with the lowest AIC. The LCT was taken as the value, in degrees C, of the breakpoint that separated the initial flat or falling leg from the first rising leg. If the preferred model by AIC had the first breakpoint as a decreasing point, we used the next-best model with an increasing initial breakpoint.

#### High-resolution respiration of isolated mitochondria

As endotherms, birds produce a substantial amount of their body heat via non-shivering thermogenesis, relying on proton leak across the inner mitochondrial membrane to dissipate the proton-motive force as radiant heat rather than usable work. Unlike mammals, birds do not appear to possess specialized thermogenic tissues such as brown fat, instead relying on expression of avian uncoupling protein (AvUCP) across tissues to uncouple mt respiration and generate heat (Evock-Clover et al., 2002; Teulier et al., 2010). Evidence supports the role of leak respiration in birds in developmental and seasonal acclimation to temperature (Correia et al., 2024; Nord et al., 2021). As such, we predicted that leak respiration would be higher in high-elevation birds and explicitly tested this assumption with a substrate-inhibitor-titration protocol on isolated mitochondria. Our protocol also allowed us to test for differences in the performance of various mitochondrial complexes, in particular Complex IV (Cytochrome C Oxidase). As the site of mitochondrial O_2_ reduction to H_2_O, Complex IV activity should be sensitive to oxygen concentration and might be the locus of adaptive differences in mitochondrial function between elevations (e.g., Scott et al., 2011).

After reaching ambient temperature at the end of the metabolic rate trial, birds had their cloacal temperature and body mass recorded again and were then euthanized by CO_2_. We did not use isofluorane for euthanasia as anaesthetics are likely mitochondrial uncouplers which might affect subsequent measurements (Kishikawa et al., 2018; Rodriguez et al., 2025). Birds were then dissected, with the brain and liver placed into ice-cold Mir05 buffer (https://www.bioblast.at/index.php/MiR05) for mitochondrial respiration. While skeletal muscle is probably the main thermogenic organ in birds (Evock-Clover et al., 2002), we were unable to assess mitochondrial function in isolated skeletal muscle preparations in the field. Instead, we examined the brain, which contributes substantially to basal metabolic rate and has moderate levels of expression of AvUCP, and the liver, which also contributes significantly to metabolic rate while having lower expression of AvUCP (Evock-Clover et al., 2002; Rolfe & Brown, 1997). A portion of the liver was also placed in RNA Later, while a sample of whole blood was placed in 100% ethanol. The heart and gut were dissected, weighed, and then dried to constant mass in a toaster oven at 100 °C. The left foot and wing were dissected, dried, and sent to the Museum of Natural History at the Universidad San Antonio Abad de Cusco (UNSAAC).

We prepared mitochondrial isolates and removed other cellular components through differential centrifugation. Brain and liver tissue were homogenized in MiR05 (https://bioblast.at/images/archive/b/bd/20241211165617%21MiR05-Kit_Manual.pdf) containing 1g/L BSA, filtered through one layer each of cheesecloth and Miracloth (Millipore-Sigma), and the resulting homogenate (1-2 mL) was centrifuged at 600 rcf for 10 min. All centrifugation was performed at 4 °C. The resulting supernatant was collected, then spun at 6000 rcf for 10 min. The supernatant was then discarded, the remaining pellets were resuspended in 1000 μL of ice-cold MiR05 without BSA, and these suspensions were centrifuged a final time at 6000 rcf for 10 minutes. The supernatant was again removed, and the pellets were resuspended in 200 μL of ice-cold MiR05 without BSA and held at 4 °C until they were used for mitochondrial respiration (10-30 mins). Total protein content of the mitochondrial isolate in Mir05 was quantified using the Qubit Broad-Spectrum protein assay, with a MiR05 blank to control for background protein content.

We then used the Oroboros Oxygraph-2K (O2K; Oroboros Instruments, Innsbruck, Austria) to quantify respiration of mt isolates using a substrate-uncoupler-inhibitor titration protocol at 41 °C (on the high end of the avian body temperature range) in a 500 μL closed chamber. We first added 500 uL of Mir05 buffer to each chamber after appropriate washes with ethanol. We calculated the oxygen flux (via electrode) of the background medium in the chamber followed by flux of the suspended mitochondrial isolate (0.25 mg protein). We then calculated the oxygen flux over stable windows (2-10 mins) in seven other respiratory states resulting from additions of the substrates and inhibitors in **Table 1**. O_2_ concentration and flux were calculated and logged every 2s in DatLab v. 7.4 (https://www.oroboros.at/index.php/product/datlab/). We analyzed mt respiration rates as flux control factors (FCFs) (Gnaiger, 2014), which are unitless metrics that normalize oxygen flux based on the addition of different substrate and inhibitors and standardize responses to 1. This has the benefit of allowing direct comparisons of respiration among samples where initial protein content or number of mitochondria added may have varied widely. In other words, by using FCFs we were able to quantify whether a particular species had a particularly high response to a certain substrate, irrespective of whether samples from that species had variable mitochondrial concentrations. We calculated 6 FCFs (**Table 2**) that correspond to proportional increases or decreases in oxygen consumption after the addition of various substrates and inhibitors. We also calculated baseline and maximum respiration value, measured in pmol O_2_ s^-1^ mL^-1^ normalized to the amount of mt protein added (**Table 2**). In our protocol, we considered maximal respiration to be the peak of the state reached after the addition of ASC+TMPD.

**Table 2.1:** Respiratory states achieved after additions of substrates and inhibitors.

| State | Addition | Quantity + Concentration added | Role | Effect on $\text{O}_2$ flux |
| --- | --- | --- | --- | --- |
| Background | MiR05 with BSA |  | Buffer | Baseline |
| Mitos | Isolated mitochondria | 0.25 mg, various concentrations |  | Small increase |
| MPG | Malate, Pyruvate, Glutamate | 1 $\mu\text{L}$ 3mM malate, 2.5 $\mu\text{L}$ 10 mM pyruvate, 2.5 $\mu\text{L}$ 10 mM glutamate | Complex I substrates | Small increase |
| ADP | ADP | 1.25 $\mu\text{L}$ 200 $\mu\text{M}$ | Complex V substrate | Big increase |
| S | Succinate | 10 $\mu\text{L}$ 5 mM | Complex II substrate | Big increase |
| Rot | Rotenone | 2.5 μL, 5 μM | Complex I inhibitor | Big decrease |
| AA | Antimycin-A | 2 μL 5 μM | Complex III inhibitor | Small decrease |
| CytC | Cytochrome C | 1 μL 25 μM | Complex IV substrate | Negligible increase |
| ASC+TMPD | Ascorbic Acid & *N,N,N',N'*-Tetramethyl-*p*-phenylenediamine dihydrochloride | 1.25 μL 10 mM ASC, 1.25 μL, 300 μM TMPD | Complex IV substrate & substrate reducer | Huge increase |

**Table 2.2:** Flux Control Factors and other metrics calculated from respiratory states in Table 1.

| FCF | Calculation | Significance |
| --- | --- | --- |
| Leak FCF | $1 - (\text{MPG}/\text{Mitos})$ | Non-phosphorylating respiration in the presence of electron-donating Complex I substrates without ADP, attributable to leak across the inner mitochondrial membrane, relative to baseline mitochondrial respiration |
| ADP FCF | $1 - (\text{ADP}/\text{MPG})$ | Phosphorylating, Complex-I linked respiration after ADP addition relative to before |
| Succinate FCF | $1 - (\text{S}/\text{ADP})$ | Phosphorylating, Complex I & II linked respiration after addition of electron-donating Complex II substrate (succinate) |
| Rotenone FCF | $1 - (\text{S}/\text{Rot})$ | Phosphorylating, Complex II linked respiration after inhibition of Complex I by rotenone |
| Antimycin A FCF | $1 - (\text{Rot}/\text{AA})$ | Phosphorylating, Complex II linked respiration after inhibition of Complexes I & III by rotenone & antimycin-A |
| ASC+TMPD FCF | 1-(ASC+TMPD/Cyt C) | Phosphorylating, Complex II & IV linked respiration after inhibition of Complexes I & III by rotenone & antimycin-A, in the presence of saturating levels of Cytochrome C |
| Baseline Respiration | Mitos/[protein] | Mitochondrial baseline respiration normalized to protein content |
| Maximal Respiration | ASC+TMPD/[protein] | Highest respiration achieved during the trial normalized to protein content |

## Results

### Selection on Mitochondrial DNA Across Elevation in Tapaculos

#### Mitonuclear discordance and independent origins of high-elevation lifestyles

MrBayes recovered similar topologies from both partition schemes. Because the tree based on the codon position-partition (Fig. 2.2) better matched the mitochondrial and nuclear results of Cadena et al. (2020), we chose to use this as the basis for our analyses of selection. Our results confirmed some patterns of mitonuclear discordance found by Cadena et al. (2020), including that the mt genome of *S. gettyae* seems to have been derived from within *S. acutirostris* and that that of *S. sanctaemartae* appears to have come from within *S. atratus*, despite reciprocal monophyly of their nuclear genomes. Additionally, we confirmed the finding that the taxa currently known as *S. altirostris*, *S. acutirostris*, and *S. latrans* are all polyphyletic with regards to the mt genome and may warrant classification into multiple distinct species. We also confirm the mitochondrial monophyly of the deeply divergent sequences assigned to *S. speluncae*, which ranges over several disjunct mountain ranges of Eastern Brazil. This degree of divergence, with the deepest splits as old as 5 mya (Cadena *et al*. 2020), is more severe than that of many distinct sister species in the genus and suggests cryptic diversity that may warrant re-classification to multiple species within what is now *S. speluncae*. Like Cadena & Céspedes, (2020), we find multiple independent origins of high-elevation living across the phylogeny of the Rhinocryptidae, giving us increased power to detect convergent selection on the mt genome.

**Figure 2.2:**
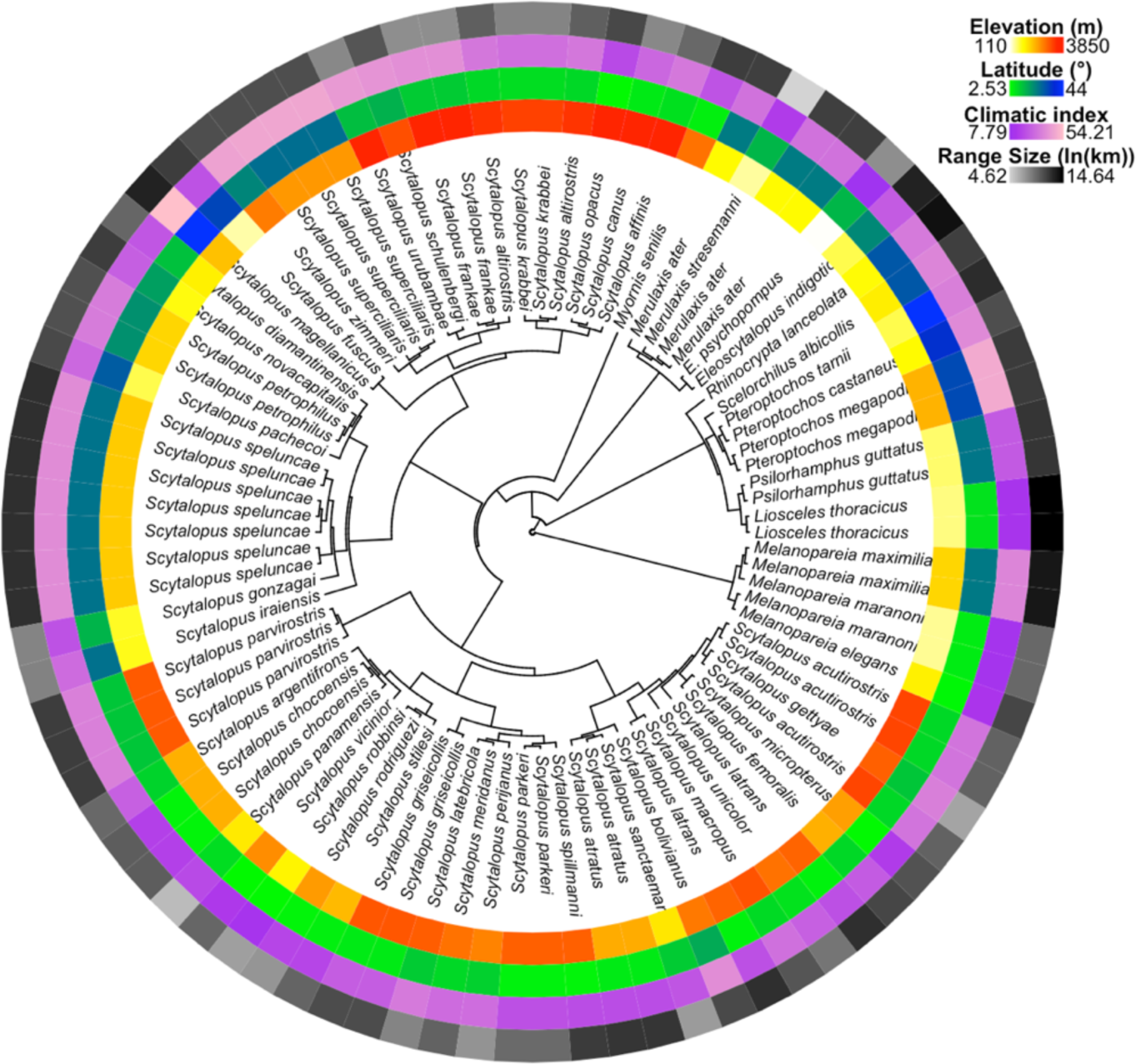
Phylogeny of the 85 species used in the study, with heat maps corresponding to the values of environmental variables calculated from IUCN shapefiles. Innermost ring: Midpoint elevation, meters above sea level; second ring: Midpoint latitude, in degrees; third ring: Climatic Index, unitless; fourth ring: Range size, log units of square kilometers. Note that taxa of high elevation and high latitude are spread throughout the phylogeny and are distinct sets from each other.

#### Positive, intensified selection on mt genes of tapaculos at the highest elevations

None of the four models binning *d*_N_/*d*_S_ estimates for species in the phylogeny (by elevation, latitude, climatic index, or range size) was preferred to a null, one-rate model by likelihood ratio test. This was expected, as these models are high in parameters, and tip bins contain little phylogenetic information relative to the background bin. However, we constrained tips to these bins explicitly to examine variation in estimated *d*_N_/*d*_S_ values among tips and contextualize results from RELAX. *d*_N_/*d*_S_ values in each bin were relativized to the null rate (*d*_N_/*d*_S_ = 0.00858) and visualized on a log scale (Fig. 2.3). *d*_N_/*d*_S_ values tended to increase with increasing midpoint elevation, with the highest values among species with midpoint elevations over 3000 m. *k*-values from RELAX also increased with elevation, largely due to species with midpoint elevations over 3000 m having *k* > 1 (*P* < 0.001, LRT), indicating positive, intensified selection on mt genes at the highest elevations. *d*_N_/*d*_S_ tended to decrease with increasing latitude, but all bins with latitude values above 20 ° failed to converge, leaving a relatively narrow range of latitudes to compare across. *k*-values tended to decrease across this range, with the bin at the lowest latitude displaying significantly relaxed positive selection relative to background (relative *d*_N_/*d*_S_ > 1, k < 1), while those at slightly higher latitudes displayed significantly relaxed purifying selection (relative *d*_N_/*d*_S_ < 1, *k* < 1). *d*_N_/*d*_S_ tended to decrease with increasing values of the Climatic Index, while *k*-values increased but remained below 1 for all bins. This suggests that species with CI ≤ 20 experienced relaxed purifying selection (relative *d*_N_/*d*_S_ > 1, *k* <1) while species with CI ≥ 25 experienced relaxed positive selection (relative *d*_N_/*d*_S_ < 1, *k* <1), with a tendency for selection to become less relaxed at higher CI values. However, the test for relaxation of selection was not significant for either the highest or lowest CI bins. *d*_N_/*d*_S_ also tended to decrease with increasing range size, but values above and below zero were found on both sides of the spectrum. All bins exhibited *k*-values below zero, with a tendency to increase with range size.

**Figure 2.3:**
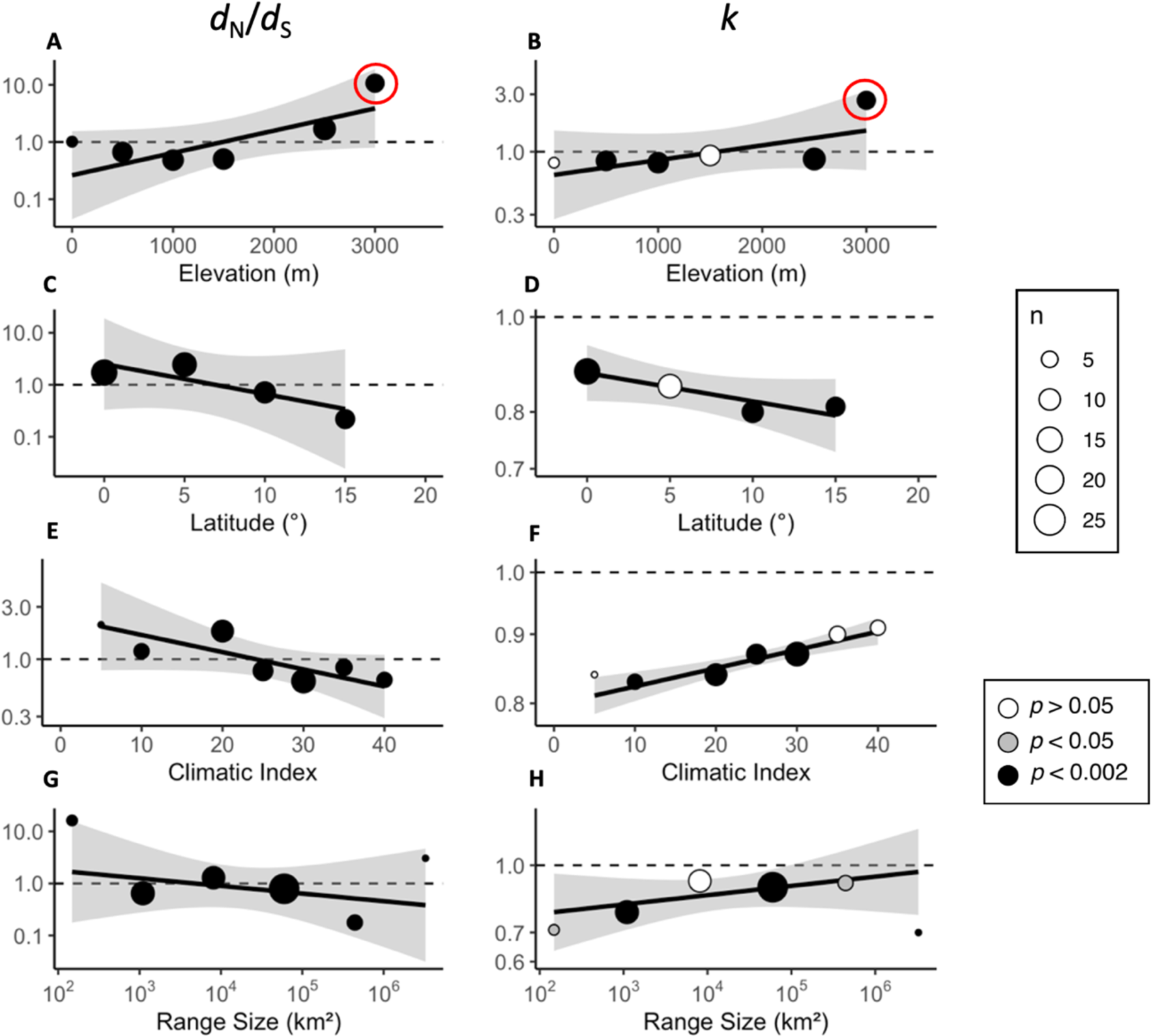
Trends in *d*_N_/*d*_S_ values and k-values across elevation for 85 species of tapaculos and allies in the families Rhinocryptidae (tapaculos) and Melanopareiidae (crescent-chests). All y-axes are on a log-scale. Black trendlines are weighted by sample size of each bin, while the gray margin represents the 95% confidence interval. Across midpoint elevations, *d*_N_/*d*_S_ (A) and k-values (B) tended to increase. Species with midpoint elevations above 3000 m (red circles) were the only ones across the dataset determined to have experienced significantly intensified positive selection (relative *d*_N_/*d*_S_ > 1 & k > 1). Across increasing midpoint latitudes, *d*_N_/*d*_S_ (C) and k-values (D) tended to decrease, with relaxed positive selection at latitudes below 10 ° and relaxed purifying selection at higher latitudes. A similar pattern was seen across the Climatic Index (E & F), with species at lower (milder) values displaying relaxed positive selection and species at higher (harsher) values displaying relaxed purifying selection. However, in this instance, k-values tended to increase with CI (F). Across variation in range sizes, *d*_N_/*d*_S_ values decreased(G) and k-values increased (H), with an idiosyncratic pattern of significantly relaxed positive and significantly relaxed purifying selection on both ends of the spectrum.

### Field Metabolic Rates and Mitochondrial Physiology Across an Elevation Transect

#### Temperatures fall with increasing elevation and exposure

For two microhabitats, both 2 m above the ground and in dense vegetation near the ground, average daily temperatures declined with increasing elevation (ANOVA; 2m: *d.f.* = 3, *F* = 142.3, *P* < 0.0001; ground: *d.f.* = 3, *F* = 354.1, *P* < 0.0001; Fig. 2.4A, B). While averages were similar for the 2 m and ground values, these microhabitats experienced different temperature ranges, with higher highs and lower lows for the exposed, 2 m loggers. Daily maximum temperatures did not decrease significantly with elevation at 2 m (ANOVA; *d.f*. = 3, *F* = 2.726, *P* = 0.0513) but did decrease with elevation at the ground level (ANOVA; *d.f.* = 3, *F* = 23.09, *P* < 0.0001). Daily minimum temperatures decreased with elevation in both microhabitats (ANOVA; 2m: *d.f.* = 3, *F* = 400.1, *P* < 0.0001; ground: *d.f.* = 3, *F* = 506.4, *P* < 0.0001).

**Figure 2.4:**
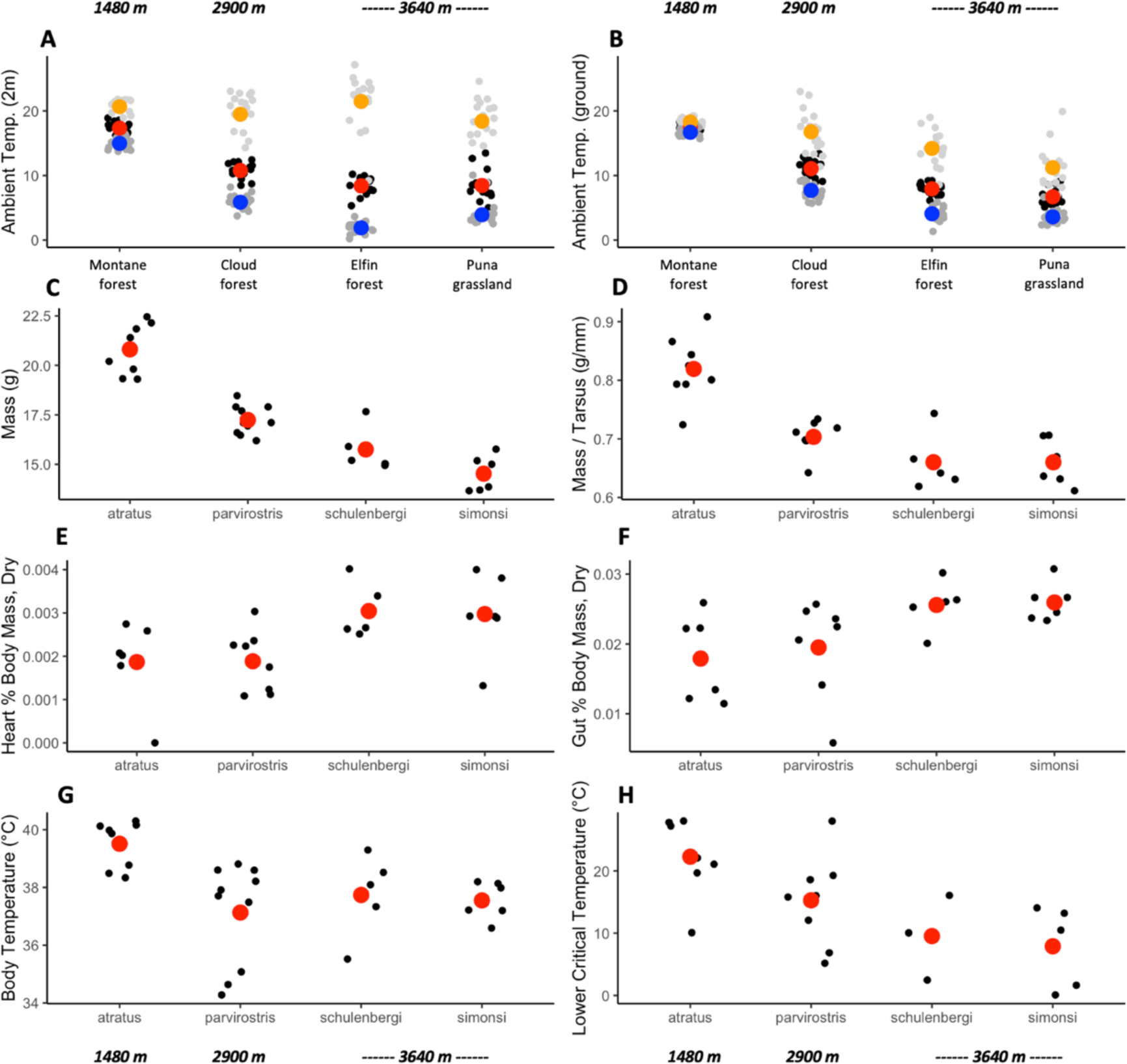
Change in environmental, morphological and physiological parameters among 4 species of *Scytalopus* replacing each other with increasing elevation. Four habitats (A&B) and four corresponding species (C-H) were compared across three elevations, which are given at the top and bottom of each column. A) Ambient temperature two meters above the ground in the habitat of each species for a common 17-day period July-August 2023 (black points = daily average temperatures, light gray points = daily maximum temperatures, dark grey points = daily minimum temperatures; red circle = average of daily average temperatures, orange circle = average of daily maximum temperatures, blue circle = average of daily minimum temperatures); B) ambient temperature within vegetation near the ground for the same period; C) Mass (g), D) Mass relative to tarsus size (g/mm), E) Dried weight of the heart as a percentage of body mass, F) dried weight of the gut as a percentage of body mass, G) Cloacal temperature (pre-trial), H) Lower Critical Temperature, based on breakpoint analysis.

#### Morphology and body temperature change with elevation in tapaculos

We captured 29 *Scytalopus* individuals (8 *S*. *atratus*, 10 *S. parvirostris*, 5 *S. schulenbergi*, and 6 *S. simonsi*) and were able to obtain both mitochondrial respiration and whole-organismal metabolic rate information for 23 of them, with the others having information for only one kind of measurement. The four species decreased in body mass significantly and in order with increasing elevation (ANOVA; *d.f.* = 3, *F* = 50.68, *P* < 0.0001, Fig. 2.4C), with the lowland *S. atratus* being the heaviest (20.81 ± 0.46 g), followed by the mid-elevation *S*. *parvirostris* (17.24 ± 0.23 g), the high-elevation *S. schulenbergi* (15.75 ± 0.51 g) and the highest-elevation, grassland-specialist *S. simonsi* (14.53 ± 0.37 g). This pattern of decreasing size with elevation was mirrored in tarsus (ANOVA; *d.f.* = 3, *F* = 21.93, *P* < 0.0001), head-bill (*d.f.* = 2, *F* = 25.35, *P* < 0.0001), and wing chord measurements (*d.f.* = 3, *F* = 10.68, *P* < 0.0005). Mass-corrected for tarsus size also decreased with elevation, with *S. atratus* being relatively heavier than the other three species (ANOVA; d.f. = 3, F = 21.51, p < 0.0001, Fig. 2.4D). Organ mass was relatively larger in the smaller species found at higher elevations, both for the gut (ANOVA; *d.f.* = 3, *F* = 3.623, *P* = 0.0299, Fig. 2.4E) and heart (*d.f.* = 3, *F* = 3.798, *P* = 0.0255, Fig. 2.4F). *S. atratus* also maintained a higher body temperature at the beginning of trials than the other three species (ANOVA, *d.f.* = 3, *F* = 5.3, *P* < 0.01, Fig. 2.4G).

### High-elevation tapaculos have lower basal metabolic rates, cooler LCTs, and higher summit metabolism

Variation in metabolic rates within each species increased with decreasing temperature (Figure 2.5), as average within-species standard deviation was 0.0103 at 28°C vs. 0.0344 at 2 °C). Our measure of basal metabolic rate (metabolic rate at 28 °C, above the LCT of all species) varied significantly among species, with the lowland and middle-elevation species having significantly lower BMR than the two highland species (ANOVA; *d.f.* = 3, *F* = 4.814, *P* = 0.0124; Fig. 2.5, black asterisk). Metabolic rate was also nearly significantly different among species at 22 °C (ANOVA; *d.f.* = 3, *F* = 3.077, *P* = 0.0556; Fig. 2.5, gray asterisk). At lower temperatures metabolic rates converged, and did not differ significantly among the four species, or among the three or two species for which estimates were available at the lowest temperatures. We also tested for differences when the two high-elevation species (which had their BMR measured at the same elevation and in the same protocol) were grouped together, and found that metabolic rate differed significantly at both 28 °C (ANOVA; *d.f.* = 2, *F* = 7.551, *P* = 0.00386) and 22 °C (*d.f.* = 2, *F* = 4.853, *P* = 0.0206) but not at lower temperatures. The difference between these two high-elevation species and medium-elevation *S. parvirostris* was nearly significant at −2 °C (ANOVA; *d.f.* = 2, *F* = 4.695, *P* = 0.0555).

**Figure 2.5:**
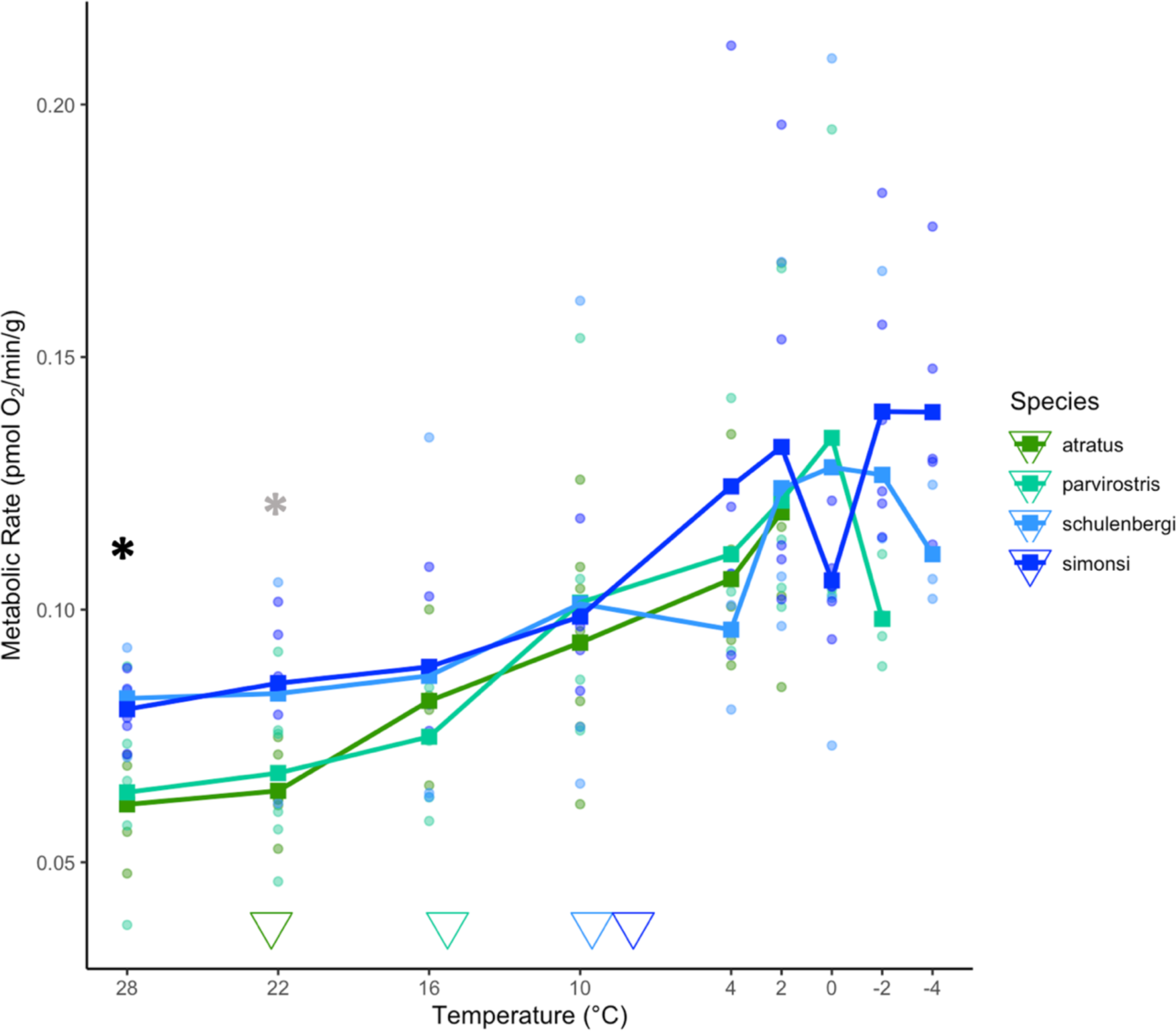
Mass-corrected metabolic rate over decreasing temperatures for four species of *Scytalopus* that replace each other with increasing elevation. Dots are individual estimates, while squares are averages of each species. Basal metabolic rate (at 28 °C) differed significantly among species by ANOVA (black asterisk), while metabolic rate at 22 °C was nearly significant (gray asterisk). Variation among all four species was not significant at colder temperatures. Triangles denote the average Lower Critical Temperature (LCT) of each species.

LCTs based on breakpoint analyses varied significantly between species (Fig. 2.4H; ANOVA; *d.f.* = 3, *F* = 5.384, *P* = 0.007), with highest average LCT in *S. atratus* (22.27 ± 2.25 °C), followed by *S. parvirostris* (15.27 ± 2.16 °C), *S. schulenbergi* (9.53 ± 3.05 °C), and *S. simonsi* (7.89 ± 2.69 °C). Summit metabolism (maximum VO_2_) was reached at −2°C for *S. simonsi*, the highest elevation species, at 0 °C for *S. schulenbergi* and *S. parvirostris*, and at 2 °C for *S. atratus*. However, because *S. atratus* could not be assayed below 2 °C, we could not determine whether its true summit metabolism had been reached. Of the three species that could be assayed below 0 °C, only *S. simonsi* (the highest species) maintained an elevated metabolic rate, reaching its highest rates at the last measurements of the trial (−2 °C and −4 °C) after a transitory decrease in rate around 0 °C. *S. parvirostris* and *S. schulenbergi*, by contrast, decreased their metabolic rates at −2 °C and −4 °C, respectively, the last measurements of their trials. *S. simonsi* had the highest summit metabolism at 0.139 ± 0.011 pmol O_2_/min/g, followed by 0.134 ± 0.031 O_2_/min/g for *S. parvirostris*, and 0.128 ± 0.041 pmol O_2_/min/g for *S. schulenbergi*. Like *S. simonsi,* both high-elevation species (*S. simonsi* and *S. schulenbergi*) appeared to experience transitory decreases in metabolic rate, at 4 °C and 0 °C, respectively, before reaching their summit metabolism.

At the end of trials, birds were weighed and had their body temperature measured again. Smaller, high-elevation species tended to lose more of their body mass during the trial, though they were also subjected to longer trials reaching lower temperatures. *S. simonsi* was the only species to increase its internal body temperature by the end of the trial, by an average of 0.45 ± 0.67 °C, while the other species all had lower body temperatures at the conclusion of the trial than their starting temperatures.

#### Mitochondrial respiration is broadly similar among low- and high-elevation tapaculos

Mitochondrial respiration rates in the liver and brain, assessed with flux control factors (FCFs), did not differ significantly among the four species in any parameters. However, because we had low sample sizes for some metrics, especially in *S. schulenbergi* and *S. simonsi*, we combined the two highest-elevation species into one category and re-examined variation. Using this three-group categorization, most metrics still did not show significant variation among groups, except for maximum respiration in the brain, with *S. atratus* having significantly higher respiration than *S. parvirostris* and the combined high-elevation species (ANOVA; *d.f.* = 2, *F* = 4.1077, *P* = 0.04375). Absolute rates, relative to protein concentration, also did not tend to differ between the brain and liver, except in peak ADP flux where activity in the brain was nearly significantly higher (ANOVA; *d.f.* = 1, *F* = 3.932, *P* = 0.0534). Respiration rates in the liver and brain were correlated at the beginning of the experiment, but tended to become uncorrelated as more substrates and inhibitors were added (Figure 2.6). For example, there were significant correlations between the brain and liver in baseline mitochondrial respiration normalized to protein content (Fig. 2.6) and in the MPG FCF (Leak respiration; Fig. 2.6B) and ADP FCF (Complex-I driven respiration; Fig. 2.6C), but not in metrics involving the response to succinate or ASC+TMPD (Fig. 2.6G).

**Figure 2.6:**
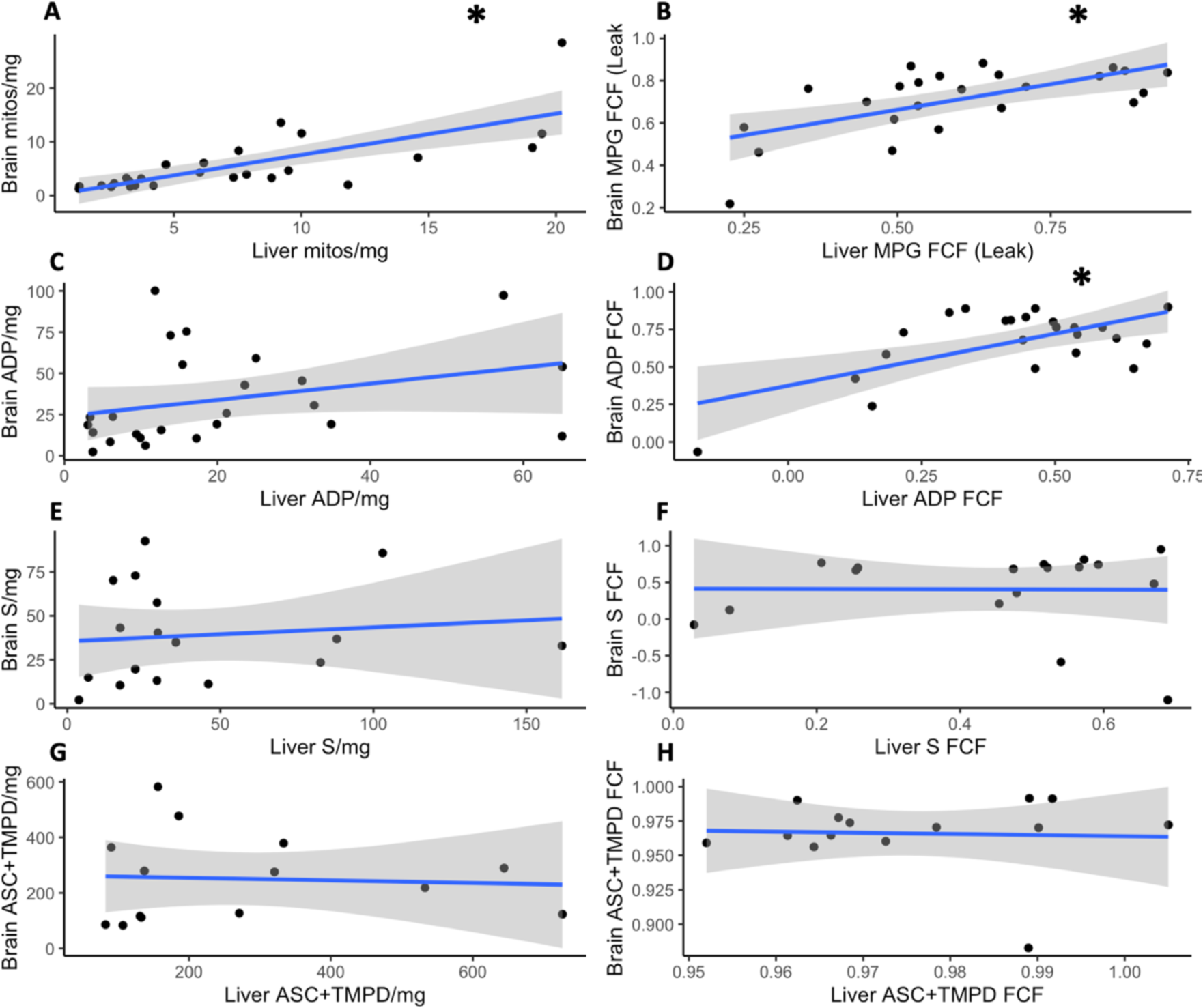
Brain and liver mitochondrial respiration variables tended to be correlated at the beginning of the experiment become uncorrelated as more substrates and inhibitors were added. The first column (A, C, E, G) depicts mt respiration states normalized to protein concentration, the second (B, D, F, H) depicts flux control factors (FCFs). Correlations in A, B, and D (black asterisks) were significant at *P* < 0.05.

## Discussion

Adaptation to elevation is expected to exert selection on the physiology of organisms that may drive genetic differentiation and potentially contribute to reproductive isolation. Here, we detected a signature of intensified positive selection in some of the core metabolic genes – the mitochondrial electron transport system proteins – in tapaculos and allies with midpoint elevations above 3000 m (Fig. 2.3A, B). Then, taking metabolic rates in the field of four species that replace each other with elevation across this threshold, we found that species at higher elevations have higher basal metabolic rates (Fig. 2.5), cooler Lower Critical Temperatures (Fig. 2.4H), and a trend towards higher summit metabolism than lower-elevation species (Fig. 2.5). However, we did not detect pervasive differences in mitochondrial function between species, finding only moderately lower levels of maximum mitochondrial respiration in high-elevation species when grouped together. While this may have functional significance, differences in basal metabolic rate are more easily attributable to the higher relative organ mass of high-elevation species (Fig. 2.4E, F). Internal organs including the heart and gut can represent up to 60% of basal energy expenditure in endotherms (Rolfe & Brown, 1997) and variation in organ size correlates with variation in BMR (Konarzewski & Diamond, 1995). Some birds display seasonal increases in organ mass and BMR in the winter (Zheng et al., 2014) or in response to experimental cold acclimation (Vézina et al., 2006). To our knowledge, this is study provides the first evidence of interspecific differences in relative organ mass that likely play a role in supporting adaptive differentiation in metabolic rates.

The question of whether high-elevation birds have generally elevated metabolic rates has been somewhat controversial. A negative relationship between BMR and ambient temperature has long been recognized for endotherms (Merchant et al., 2024; White et al., 2007), which allows them to maintain remarkably consistent internal temperatures across a wide range of environments. However, studies of bird communities that span elevation gradients in New Guinea (McNab, 2013) and Peru (Gutierrez-Pinto et al., 2021) have failed to find a general pattern of elevated metabolic rates for high-elevation birds. Surprisingly, to our knowledge only one study has directly examined paired high and low elevation congeners, finding higher basal metabolic rate in most high-elevation species (Gutierrez-Pinto et al., 2021). One complicating factor in the relationship between elevation and BMR is that, while low temperatures are associated with higher BMR, lower oxygen levels are associated with lower BMR (Gu & Jun, 2018; Lu et al., 1999; Nathaniel et al., 2012). While low BMR is a documented plastic response to hypoxia within individuals, it is not clear whether the same response is fixed by selection for life under hypoxic conditions. BMR is partly heritable (average *h*^2^ = 0.19) and appears to be more heritable in endotherms (Pettersen et al., 2018), perhaps because ambient temperature exerts its impact indirectly. In deer mice, highland natives have higher maximal metabolic rates under simulated hypoxic conditions, demonstrating an impact of local adaptation to high altitudes (Bautista et al., 2024; Cheviron et al., 2013). In this species, highland genetic ancestry has a bigger influence on metabolic rate than acclimation by lowlanders. Thus, while lower metabolic rates might be expected in response to acute hypoxia, the present study complements existing evidence that high-altitude endotherms instead evolve higher aerobic capacities for increased heat-production despite the challenge of low oxygen availability. Because we could not control for oxygen concentration in our experiments, these four species experienced the same cold challenge at progressively lower oxygen concentrations. The impressive ability of the higher-elevation species to maintain the same or greater oxygen consumption under these more hypoxic conditions suggests an increased efficiency of oxygen use that may have a genetic basis. However, we cannot disentangle acclimation and adaptation effects in the present experimental design.

### Physiological Basis for Differences in Metabolic Rate

We hypothesized that metabolic differences between species might be downstream consequences of differences in mitochondrial function, such as in LEAK respiration (representing the thermogenic state of the inner mitochondrial membrane) or the ADP and S FCFs (representing phosphorylating respiration driven by different complexes in the presence of saturating substrates). However, contrary to our predictions, we detected few significant differences between species in the measures of mitochondrial function we examined in either the brain or liver. The one significant difference among our data suggested that the lowland *S. atratus* had higher maximal respiration than the other species in the brain, with a similar but non-significant trend in the liver. We caution the interpretation of this result, given the low sample sizes and high variability within species; however, one possibility is that *S. atratus* has higher activity of Complex IV (cytochrome C oxidase) in the brain. The design of our protocol meant that maximal respiration at the end of the experiment (in response to ascorbic acid and TMPD) was carried out after Complex I and III had been deactivated, and saturating concentrations of Cytochrome C were provided. Because Complex II activity (respiration in the presence of succinate, after Complex I was deactivated by rotenone) did not differ between species, the difference in maximal respiration could instead correspond to a difference in Complex IV activity. Complex IV is the site of most mitochondrial oxygen consumption in the electron transport system, and decreased Complex IV activity has been reported as a response to high-altitude exposure (Howald et al., 1990; Levett et al., 2012). It is possible that decreased Complex IV activity in our high-elevation species relative to *S. atratus* is an evolved response to hypoxia, or it may reflect phenotypic flexibility that all species are capable of under hypoxia.

Regardless of the role that Complex IV activity may play in metabolic differences, the overall similarity in mitochondrial functional profiles suggests that much of the differentiation in whole-organism metabolic rates and gross morphology observed here is driven by variation at other levels of organization. Metabolic rate is a highly integrative trait dependent not just on enzyme activities and mitochondrial oxygen consumption, but on the density and distribution of mitochondria, the volume and organization of mitochondria-rich tissues, and the regulation of tissue-level metabolism by vasoconstriction and dilation, heart rate and respiration rate, and changes in heat conductance across the skin (Iverson et al., 2020). For instance, we observed significant differences in organ mass (heart and intestine) that could support higher basal metabolic rates relative to body size in the two high-elevation species, even if overall mitochondrial respiration were similar between them. Larger organs will also support higher summit metabolism, and there may be other differences in characteristics of the circulatory system we did not measure. Additionally, by maintaining lower body temperatures (Fig. 2.4G), high-altitude species can reduce the overall metabolic demand on these tissues (Freeman et al., 2022; Wearing & Scott, 2022). While we did not detect substantial differences in mitochondrial function between species, we assayed mitochondrial function under idealized conditions. For instance, isolated mitochondria are not constrained by oxygen delivery from the circulatory system and thus all assays took place under presumably saturating conditions, even at high elevation. Additionally, we are unable to replicate the cellular environment induced by thermogenic demand, in which nuclear-encoded uncoupling proteins (eg., AvUCP) are likely to be upregulated to drive heat production across the inner mitochondrial membrane. Hypoxia and thermogenic demand are stressors that might exacerbate functional differences among species at the mitochondrial level largely through changes in the nuclear regulatory environment; thus, there may be other genetically-encoded differences in mitochondrial function that we could not detect.

### Allometry and Competition Among Species

The higher proportional organ masses we observed in high-elevation species stand out because these species are also smaller, both absolutely and in in their weight relative to their tarsus size (Fig. 2.4C, D). This contravenes traditional predictions from Bergman’s rule (that cold-adapted species should be larger to reduce their surface area to volume ratio) and Allen’s rule (that cold-adapted species should have shorter extremities relative to their body size). However, a recent analysis of birds found that in small species, tarsus length is negatively related to temperature due to unknown mechanisms; additionally, in relatively long-legged birds, body size increases with temperature (Frӧhlich et al., 2023). These contextual inversions of traditional temperature-size rules accurately describe the pattern we find among the small, long-legged tapaculos.

Whatever ecological constraints drive tapaculo species at higher elevations to be both smaller and relatively longer limbed, they appear to have compensated for any increased heat loss by maintaining larger internal organs that support higher basal metabolism.

The pattern observed here across all four species fits a recently proposed paradigm for ecological replacements, the Competitive Exclusion-Tolerance Rule (Martin & Ghalambor, 2023). Supported by experiments in both animals and plants along environmental gradients, the rule suggests that species which are weaker competitors (typically, the smaller congener) are confined to the harsher part of an environmental gradient, to which they are more tolerant than the better competitor. When the species on the benign part of a gradient is removed, these smaller, less dominant species expand their range because they are tolerant of more benign conditions; however, when the species removed is from the harsh part of the gradient, the stronger competitor does not expand its range, or expands less, because it is limited by its environmental tolerance.

Essentially, the Competitive-Exclusion-Tolerance Rule formalizes with data a theory dating back to Darwin (1859) that species are constrained by biotic factors on one edge of their range and by abiotic factors on the other. The four *Scytalopus* species here illustrate this pattern well – each species at progressively higher elevations is smaller and, presumably, a poorer competitor, but each shows evidence of progressively greater cold tolerance. This chain reaction of competitive exclusion suggests that with global warming, each species will be outcompeted on its warm edge (rather than being intolerant of warmer conditions) and will expand on its cold edge by competitively excluding the next higher species. One testable prediction for the future from this hypothesis is that Upper Critical Temperatures (UCTs) may not differ much between species, or may have little connection to relevant environmental variation, because they are not under direct selection like LCTs. Daily minimum temperatures, which are rising faster than maxima in some areas due to climate change (Gil-Alana, 2018), would release larger species from environmental exclusion and may be an essential variable to monitor in mountains for understanding species upward migrations.

### Distribution on Environmental Gradients

Prior work in other organisms along the same elevation gradient (Manu Biosphere Reserve/Southeastern Peru) has established two ecological transitions with broad significance for ecological replacement and climate tracking. First, between 1500 and 1750 meters occurs the formation of the cloud base, with near-daily inundation by atmospheric moisture above this point leading to marked differences in vegetation structure and species composition (Los et al., 2021; Terborgh, 1971). Second, around 3000-4000 m occurs the forest-grassland transition (or treeline), a complex and variable ecotone formed by the interplay of climate, fire, soils, and grazing (Bush et al., 2015, 2022; Rehm et al., 2015; Rehm & Feeley, 2015; Sarmiento & Frolich, 2002). While the cloud base is rising in elevation with warming, the treeline has remained stable or even become lower in many parts of the Andes due to human impacts, primarily fire (Los et al., 2021; Rehm & Feeley, 2015, 2016). The high metabolism of the two high-elevation species on either side of the treeline ecotone (*S. schulenbergi* in the elfin forest and *S. simonsi* in the puna grasslands) and the intermediate character of the middle-elevation *S. parvirostris* in some traits suggests that the continuous decrease in temperature and oxygen availability with elevation may be what governs turnover in tapaculos. This makes sense in the context of findings from the region that turnover in birds appears to have more to do with the climatic gradient than to ecotones (Freeman et al., 2018; Terborgh, 1971, 1977). As such, a rising cloud base *per se* would not threaten birds or impact their upward migration but would merely be symptomatic of a changing climate which directly affects them physiologically. However, the treeline ecotone is more likely to represent an important physiological divide for birds; our loggers in the elfin forest and puna grasslands at the same elevation highlighted subtle differences between them, including a wider temperature range 2 m above the ground in the elfin forest, and lower maximum and average daily temperatures near the ground in the puna.

In our phylogenetic analysis, the species with midpoint elevations above 3000 m (*S. affinis, S. canus, S. frankeae, S. opacus,* and *S. urubambae*) in which positive selection on mitochondrial genes was detectable tend to occur at or above the treeline. Species in the next lowest bin, with midpoint elevations above 2500 m, live in upper cloud forest and elfin forest near or below treeline. This group was the only other elevation bin to have a relative *d*_N_/*d*_S_ value above 1, although selection was found to be relaxed like in all remaining bins. Despite their similarity to each other relative to the two lower-elevation species, *S. schulenbergi* and *S. simonsi* also seem to have metabolic differences in some of the traits that might relate to a harsher microclimate above tree-line: *S. simonsi* tended to have a lower LCT, reached a higher summit metabolism at a lower temperature, and maintained high respiration until the end of the trial without a noticeable falling leg. At the end of trials, *S. simonsi* was also the only species with a higher body temperature than when the trial began. This remarkable thermogenic capacity in a grassland species, relative to a congener captured and measured at the same elevations but from a forest microhabitat, suggests important differences in the conditions to which the two species have adapted and implies that upward migration of *S. schulenbergi* may be constrained by the availability of the forest microclimate. Because Andean treelines are failing to rise with climate change due to anthropogenic disturbance at high-elevations, *S. schulenbergi* and other elfin forest endemics may be the most vulnerable species on the gradient, as they are squeezed by competition from forest-adapted congeners from below. A reduction in anthropogenic disturbance would allow elfin forest habitat to migrate upslope, preventing reduced ranges and potential extinction as a series of migrating species encounter a hard barrier.

## Conclusion

We demonstrate here that in a clade of wide-ranging neotropical birds, species living above 3000 m display converging signals of intensified positive selection on mitochondrial genes that are not found in relation to latitude, relative climatic index, or range size. By performing experimental whole-animal respirometry in the field on four species of *Scytalopus* tapaculos that replace each other with elevation, we show that higher-living species have higher BMR, lower LCTs, and higher summit metabolism despite being smaller overall. However, experimental mitochondrial respirometry of the same birds does not support a major role for baseline differences in mitochondrial function in driving these differences in whole-organism metabolic rates; rather, high metabolic performance in the smaller, high-elevation species is likely supported by their larger internal organ masses relative to their body size. The juxtaposition of decreasing body size and increasing cold tolerance with increasing elevation suggests a trade-off between competitiveness and environmental adaptation, with competition constraining the warm range margin and tolerance constraining the cold margin. All species may be moving upward as minimum temperatures rise due to climate change, but the upward migration of *S. schulenbergi*, which inhabits the highest forested habitats, could be constrained by the stability of the Andean treeline in the face of anthropogenic disturbance. This work illustrates the complicated interplay between physiology, interspecific interactions, and human land use change in identifying species vulnerable to climate change.

## Acknowledgments

We wish to thank the staff of the Peruvian agencies SERNANP and SERFOR for facilitating permits and access, in particular Beto Pacheco, formerly of SERNANP. We also thank the staff of the Cock-of-the-Rock Lodge, including Walter Borda, and the staff of Wayqecha Biological Station, including Ruthmery Pilco Huarcaya, for support and assistance. We thank Kendra Zwonitzer for logistical support.

